# Depth of sedation with dexmedetomidine modulates cortical excitability non-linearly

**DOI:** 10.1101/2021.06.04.447060

**Authors:** Paolo Cardone, Olivier Bodart, Murielle Kirsch, Julien Sanfilippo, Alessandra Virgillito, Charlotte Martial, Jessica Simon, Sarah Wannez, Robert D. Sanders, Steven Laureys, Marcello Massimini, Vincent Bonhomme, Olivia Gosseries

## Abstract

**Background:** Cortical excitability changes across conscious states, being higher in unconsciousness compared to normal wakefulness. Anaesthesia offers controlled manipulation to investigate conscious processes and underlying brain dynamics. Among commonly used anaesthetic agents, dexmedetomidine (DEX) effects are not completely known. In this study, we investigated cortical excitability as a function of DEX sedation depth.

**Methods:** Transcranial magnetic stimulation coupled with electroencephalography was recorded in 20 healthy subjects undergoing DEX sedation in four conditions (baseline, light sedation, deep sedation, recovery). Frontal and parietal cortices were stimulated using a neuronavigation system. Cortical excitability was inferred by slope, amplitude, positive and negative peak latencies of the first component (0-30 ms) of the TMS-evoked potential. Four Generalized Linear Mixed Models (GLMM) were used to test the effect of condition and brain region over cortical excitability.

**Results:** Dexmedetomidine modulated amplitude (P<0.001), slope (P=0.0001) and positive peak (P=0.042), while the targeted brain region affected amplitude (P<0.001), slope (P<0.001), and negative peak (P=0.001). The interaction between dexmedetomidine and region had an effect over amplitude (P=0.004), and slope (P=0.009) such that cortical excitability was higher during all conditions where DEX was present as compared to the baseline.

**Conclusions:** Cortical excitability changes non-linearly as a function of the depth of DEX sedation, with a paradoxical non dose-dependent increase. The effect is region-specific, being present in the frontal but not in the parietal region. Future research should extend the current results with other anaesthetics to better understand the link between cortical excitability and depth of sedation.

## Introduction

Anaesthesia offers a unique medium to unveil consciousness mechanisms, modulating reversibly different aspects of consciousness states, depending on the nature of the drug and its dosage (for a recent review, see^1^). When an anaesthetic agent leads to an alteration of consciousness, it impacts the brain functioning in its complexity^2^, connectivity^3^, and frequency range^4^. After regaining consciousness, people might experience emergence agitation, postoperative delirium, a cognitive disorder characterised by anxiety, cognitive alterations, and/or hypo- or hyperactivity.^5^

Dexmedetomidine (DEX) is an α_2_-adrenoceptor agonist that has the potential of reducing the incidence of emergence agitation^6^ and postoperative delirium compared to other anaesthetic agents^7^. The reasons for these phenomena are still unclear. The anxiolytic, analgesic and opioid sparing properties of the molecule, as well as the absence of anticholinergic effects, improvement of sleep quality, and eventually attenuation of postoperative inflammation have been advocated to explain the reduction in the incidence of postoperative delirium^8^. Moreover, a possible quicker transition between brain states and quicker restoration of cortical communication might explain the positive effect on emergence agitation. Through its inhibiting effect on the locus coeruleus, DEX reduces the inhibition of the ventrolateral preoptic nucleus (VLPO) of the hypothalamus, which in turn exerts GABAergic inhibition of cortical arousal nuclei. This effect on subcortical sleep systems promotes a state similar to stage 2/3 non-REM sleep.^9–11^ After DEX intake, cortical and subcortical regions glucose consumption decreases, which correlates with the functional connectivity impairment in intrinsic consciousness networks, as well as between the thalamus and cortical regions within those networks.^11 12^ Network topology is also modified by DEX.^13^ Interestingly, the cortico-cortical connectivity remains partially preserved during deep sedation.^12^ This asymmetry between cortical and subcortical regions might account for partially preserved semantic processing of incoming stimuli after the loss of responsiveness, as indexed by electroencephalography (EEG).^14^ Also, functional connectivity between the thalamus and key structures of arousal and saliency detection networks is relatively preserved during DEX-induced deep sedation, which may explain the ability to rapidly restore responsiveness by vigorous external stimulation. Thus, responsiveness and information processing are modulated by DEX-induced modifications in brain activity. Finally, DEX drives a shift towards slow-wave oscillation^15 16^, while high-frequencies power (i.e., beta) can accurately predict responsiveness upon behavioural assessment^17^. These findings pave the way to investigate the link between responsiveness, depth of sedation, and relative cortical modulation.

Transcranial magnetic stimulation coupled with high-density electroencephalography (TMS-hdEEG) assesses brain response with a no-task paradigm, bypassing sensory cortices. TMS-hdEEG is a non-invasive neurostimulation technique that perturbs the brain through a local and fast change of the magnetic field. This change induces an electrical current that mimics physiological activity, leading to an endogenous-like response to the pulse. TMS-evoked potential (TEP), the averaged EEG response to the TMS pulse, captures the neural response.^2 3^ TEP at the nearest electrode to the stimulation side provides information on the local modulation of the TMS. We can operationally define cortical excitability as the amplitude, slope, and positive/negative response latencies of the first component (0-30 ms), although we remain blinded to the underlying neuronal events. Cortical excitability as measured this way is modulated by conscious states^18^, circadian rhythms, sleep, and sleep deprivation^19 20^. It also increases during unresponsive states such as NREM sleep^21^ and attentional lapses^22^, standing as a promising method to investigate reactiveness of the cortex in time and space as a function of conscious states.

In this study, we aimed to directly inquire DEX effects over cortical excitability during different levels of sedation [namely no sedation (baseline), light sedation, absence of volitional response to command (deep sedation), and recovery of volitional response (recovery)]. Following the effects described in sleep, we expected cortical excitability to proportionally increase with depth of sedation. We hypothesised that cortical excitability would be the highest during deep sedation, while there would be virtually no difference between baseline and the recovery condition after DEX intake, where subjects show behavioural responsiveness.

## Methods

### Participants

*A priori* power analysis and sample size estimation were difficult given the scarcity of research on TEPs, anaesthesia and cortical excitability. We aimed to include at least 20 subjects, as this is in the range of most TMS-hdEEG studies.^2 22^ Considering drop-out and possible technical problems, we recruited thirty healthy subjects on the university campus between February 2015 and May 2016. Participants were screened by a senior anaesthesiologist (VB) to control for the absence of any contraindications to DEX sedation, TMS, and MRI. We recruited adult healthy volunteers on the university campus with the following inclusion criteria: more than 18 years, absence of prior neurological, neurosurgical, or psychiatric history, no history of adverse events during anaesthesia or previous exposure to dexmedetomidine, no active chronic illness or medication, no contra-indication to MRI, and no ongoing pregnancy for female participants (efficient contraception or negative pregnancy test required before inclusion). Five participants were dismissed because artefact-free TEPs could not be obtained reliably during normal wakefulness, two lost interest in the study, and two were dropped for technical or logistical reasons. One subject had a minor adverse reaction to DEX infusion (pruritus without a rash or any other symptoms or signs), for which the experiment was aborted. Twenty subjects completed the entire experiment (see **Table 1**). All subjects gave their written informed consent. The study was approved by the Ethics Committee of the University and University Hospital of Liège, Belgium (number B707201422895, professor V. Seutin).

**Table 1:**
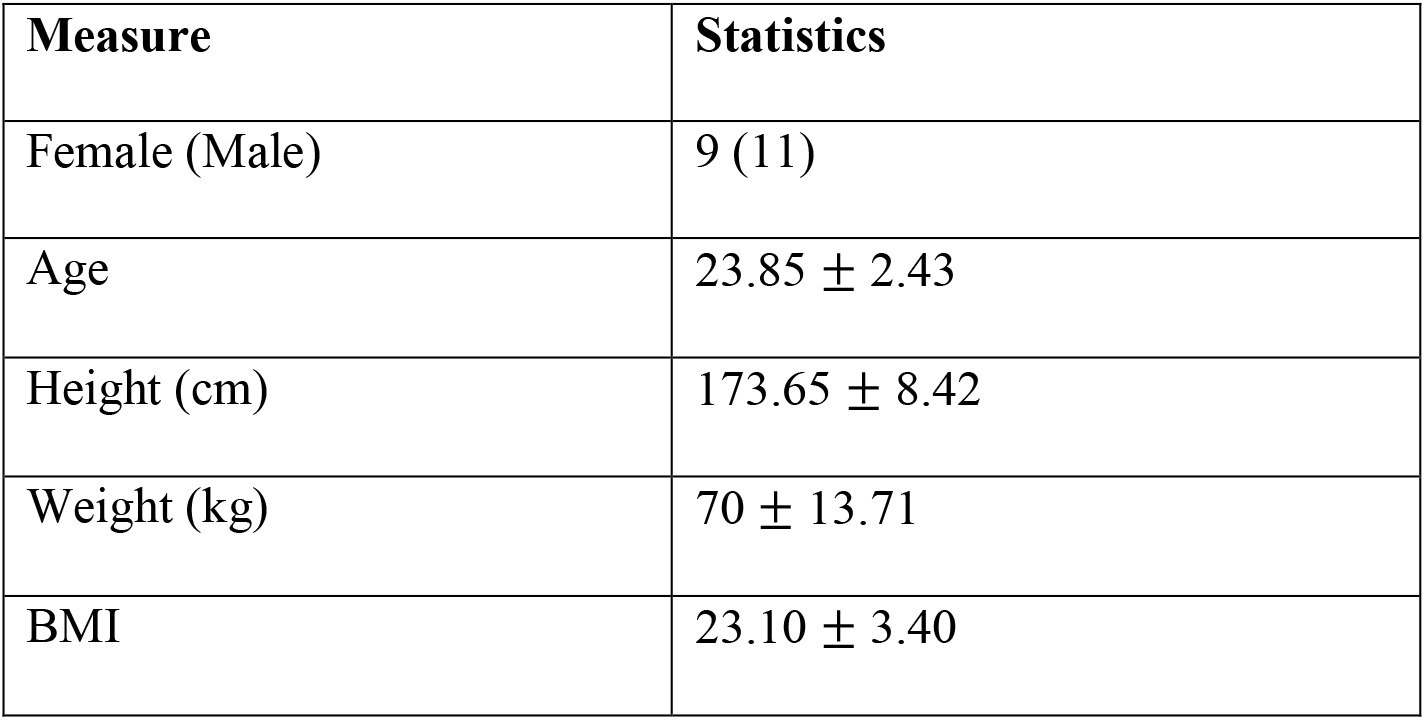
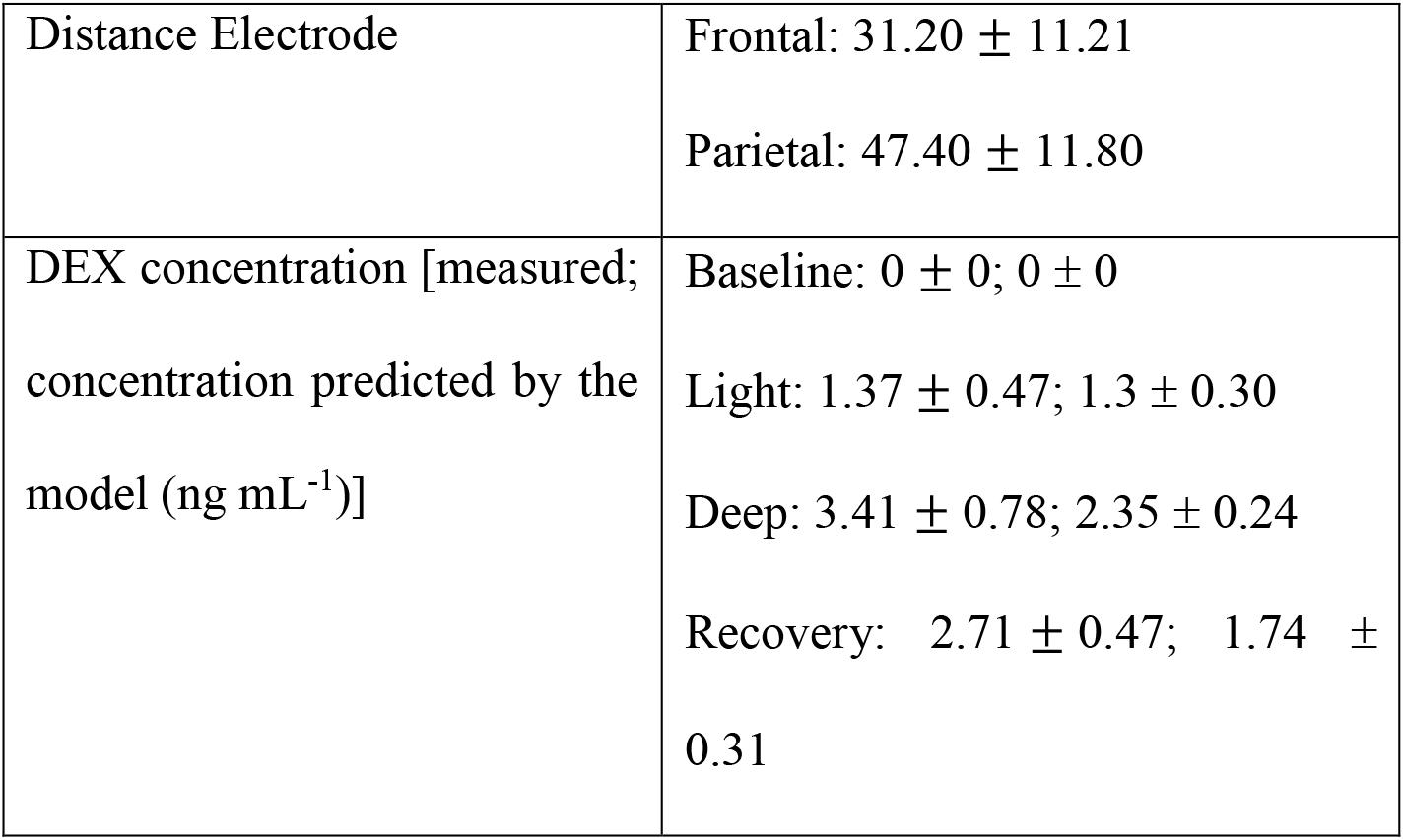
Demographics and descriptive statistics of variables. Where continuous variable, we present mean and standard deviation (mean ± SD); where categorical, we show the count for each.

| Measure | Statistics |
| --- | --- |
| Female (Male) | 9 (11) |
| Age | $23.85 \pm 2.43$ |
| Height (cm) | $173.65 \pm 8.42$ |
| Weight (kg) | $70 \pm 13.71$ |
| BMI | $23.10 \pm 3.40$ |
| Distance Electrode | Frontal: $31.20 \pm 11.21$<br>Parietal: $47.40 \pm 11.80$ |
| DEX concentration [measured; concentration predicted by the model ($\text{ng mL}^{-1}$)] | Baseline: $0 \pm 0$; $0 \pm 0$<br>Light: $1.37 \pm 0.47$; $1.3 \pm 0.30$<br>Deep: $3.41 \pm 0.78$; $2.35 \pm 0.24$<br>Recovery: $2.71 \pm 0.47$; $1.74 \pm 0.31$ |

### Experimental protocol

A visual summary of the protocol can be found in **Figure 1**. After a first screening, eligible participants underwent an MRI and a TMS-hdEEG pretest during normal wakefulness to find the most suitable brain target under stimulation of the superior parietal (Precuneus - Brodmann area 7) and premotor region (Brodmann area 6) at the midline. These brain targets were set for the experimental phase using neuronavigation (Nexstim, Helsinki. Finland). During the experiment, subjects lied on their back while venous access was installed to infuse the drug. DEX was administered intravenously using a target-controlled infusion device (TCI, height-adjusted model of Dyck^23^), providing a constant estimation of DEX plasma concentration. DEX target concentration was changed by steps of 0.5 ng mL^−1^ to achieve the desired behavioural state. Once attained, a 5-minute equilibration period without any change in target concentration allowed equilibration of concentrations between pharmacokinetic compartments, and a blood sample was drawn immediately before and after data acquisition for off-line DEX plasma concentration measurement via high performance liquid chromatography-mass spectrometry, or HPLC-MS (see Appendix). The behavioural assessment of depth of sedation was performed at the same times using the University of Michigan Sedation Scale (UMSS)^24^ and Ramsay Scale^25^. There were four conditions for each subject: “baseline”, before DEX administration; light sedation, marked by drowsiness; deep sedation, characterised by no behavioural response; recovery, with regaining in response. During the whole study, physiological parameters were monitored (ECG, peripheral blood oxygen saturation by pulse oximetry, and end-tidal CO2 levels). After a baseline TMS-hdEEG recording, DEX was increased to reach drowsiness. A 5-minute break allows concentration to stabilise, reaching light sedation, during which subjects were still able to follow a command. The level of DEX was then incremented by 0.5 ng mL^−1^ steps to induce unresponsiveness, alias deep sedation. For security reason, we did not exceed 2.5 ng mL^−1^. Lastly, the DEX concentration was decreased by 0.5 ng mL^−1^ steps to regain responsiveness to command, which was referred as the recovery condition. Once responsiveness had returned, the attained concentration was maintained constant for the duration of recordings.

**Figure 1:**
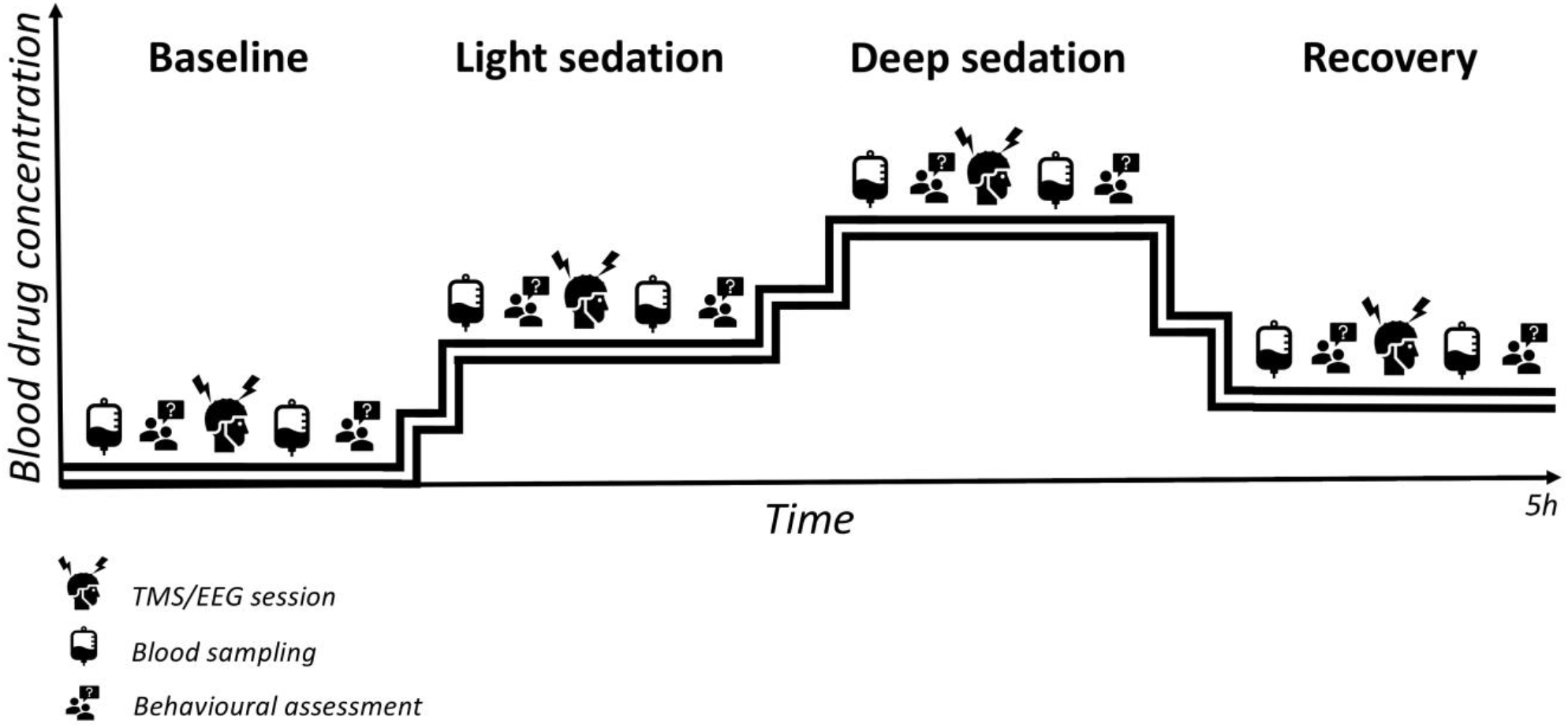
Diagram of the protocol plotted over time (x-axis, arbitrary scale) and DEX concentration (y-axis, arbitrary scale). Four conditions were set (Baseline, Light Sedation, Deep Sedation, Recovery) based on behavioural assessment. TMS-hdEEG sessions over the parietal and frontal regions were performed in each stable condition, for a total of 8 sessions per subject.

### Data acquisition

#### Magnetic resonance imaging

High-resolution structural MRI was performed on a 3-Tesla MR scanner (Allegra Prisma, Siemens, 3D isometric 1×1×1mm T1) during wakefulness, on pretesting day, just before the TMS-hdEEG session. For each participant, diffusion-weighted imaging data was acquired (not used in the study). T1 was used to perform TMS neuronavigation on the individual cortex.

#### TMS-hdEEG

A focal bipulse 8-coil (Nexstim, Helsinki, Finland) with a 3D infrared tracking position sensor was used to perform TMS delivery. Neuronavigation was implemented using glasses head tracker and the Navigated Brain Stimulation (NBS) system (Nexstim Ltd., Helsinki, Finland) that uses T1-weighted structural MR images to set stimulation target. A 64-channel TMS-compatible EEG amplifier (Eximia, Helsinki, Finland), equipped with a sample-and-hold circuit to provide TMS-artefact-free data from 5 ms post-stimulation, was used to record concurrent EEG data during TMS stimulation. Electro-oculogram (EOG) was recorded with two bipolar electrodes. EEG signal was band-pass filtered between 0.1 and 500 Hz and sampled at 1450 Hz. Prior to each recording session, electrodes impedance was set below 5 kΩ. Stimulation target and intensity were set during the pretest and were kept constant across all conditions. Left premotor and left parietal cortices were targeted and the stimulation target was chosen if there was a good TEP with no artefact. The intensity was adjusted individually to get a good signal-to-noise ratio, with an evoked electric field intensity at the cortical surface between 100 and 150 Vm^−1^. Each condition had between 200 and 250 trials, with a frequency of 0.5 Hz and a jitter of ±200 ms. A thin foam layer under the TMS coil and white noise mask were used to minimize somatosensory stimulation and auditory evoked potentials caused by the TMS click, respectively.

#### Behavioural assessment

Behavioural assessment of depth of sedation was performed using the UMSS^24^ and Ramsay Scale^25^. The four conditions had different behavioural profiles: baseline, previous to the DEX administration, was marked by a clear command-following to the verbal request ‘squeeze my hand’ (Ramsay score 2, UMSS 0); light sedation was marked by drowsiness (Ramsay score 3-4, UMSS 1-2); deep sedation was characterised by no behavioural response to any verbal command (Ramsay score 6, UMSS 4); recovery was distinguished by regaining in response after deep sedation (Ramsay score 3-4, UMSS 1-2). To exclude possible automatic response to command, other minor attentional and memory tasks were performed. These tasks included predetermined questions about simple subtractions and autobiographical memory recalls (not analysed here).

#### Blood sampling

Before and after each session we took a blood sample to calculate the real plasmatic DEX concentration. The sample was anonymized and stored at −20°C before being analysed by Orion Pharma. For more information about the blood sampling and analysis, see Appendix.

### Data analysis

#### Preprocessing of TMS-hdEEG data

Data were analysed using MATLAB (The Mathworks Inc., Natick, MA). Trial rejection was performed manually with SSP (SiSyphus Project) to eliminate trials with magnetic artefacts or ocular/muscular movements. Channels with a high level of noise were rejected. A first 1 Hz high pass filter was applied to continuous data to eliminate slow oscillating noise. Afterwards data were downsampled to 1000 Hz, then lowpassed to 80 Hz. Data were subsequently epoched from −100 to 300 ms post-stimulation. A baseline correction between −100 and −1.5 ms was applied. Trials were then averaged, using robust averaging method, to minimize noise. For more details, see previous publications where the same methods were applied.^19 20 22^

#### Cortical excitability computation

Cortical excitability was inferred from the amplitude, the slope, the positive and negative latency of the first component of the TEP, between 0 and 30 ms post-TMS. The TEP was extracted at the closest electrode to the stimulation point that did not present any artefact (distance of the electrode from the hotspot, mean ± SD, 39.29 ± 14 mm). The latency of the negative peak is the time delay between the stimulation (t_N_) and the moment at which the TEP is minimum, and ranges between 9 and 15 ms, while the latency of the positive peak (t_P_) is the time delay between the stimulation and the moment at which the TEP is maximum, and ranges between 10 and 30 ms. The amplitude refers to the peak-to-peak amplitude 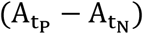, which is the microvolt change between peak 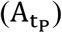 and trough 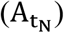, while the slope is the maximum change of the first component between t_P_ and t_N_. More details can be found in previous works.^19 20 22^ For a visual intuition, see **Figure 2**.

**Figure 2:**
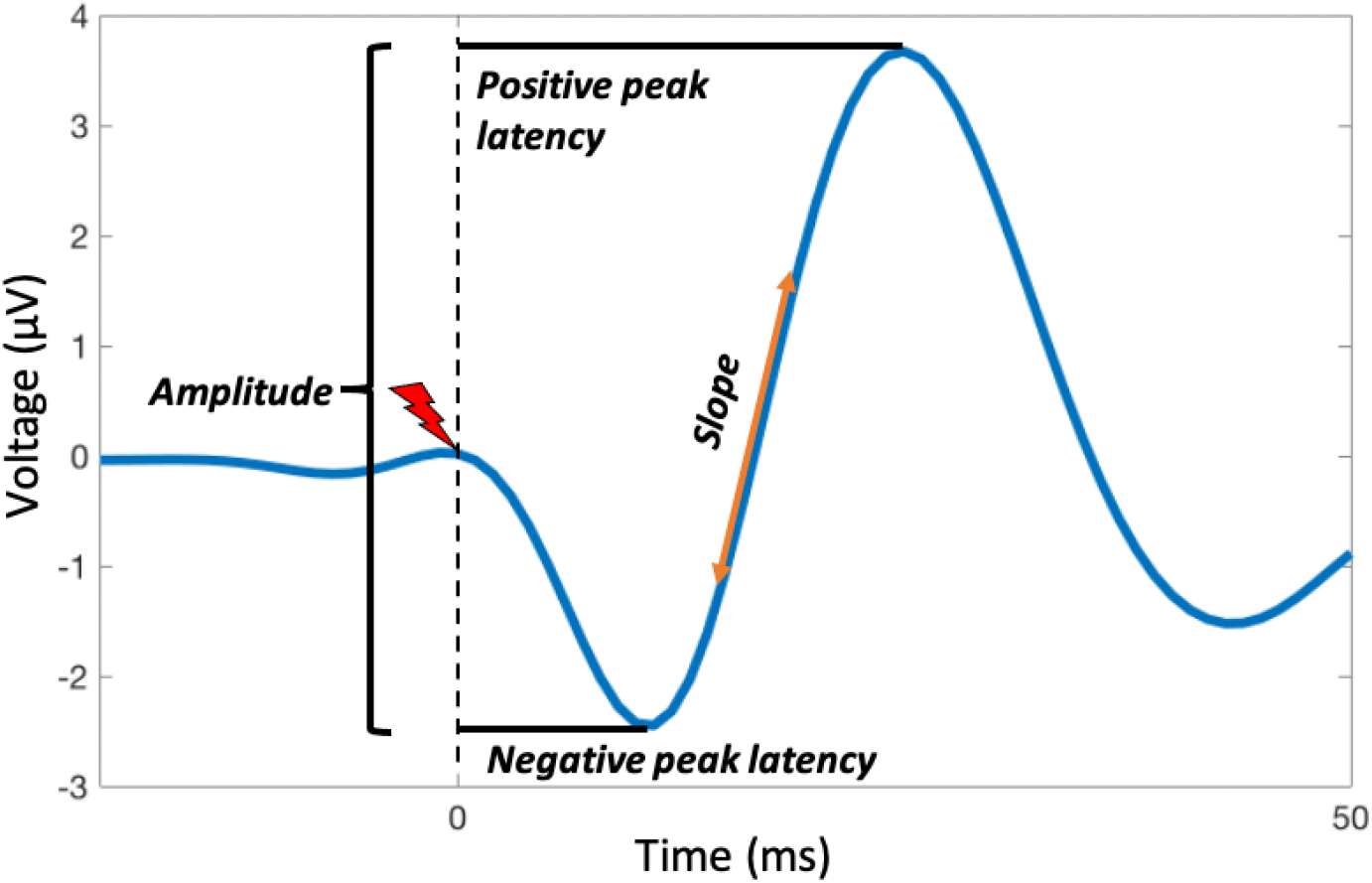
Measures of cortical excitability in the TEP (average TMS-hdEEG responses over trials). The red flash indicates the TMS pulse. We measured the peak-to-peak amplitude of the TEP in μV (here, around 6 μV), the latency in milliseconds of the negative peak (here, around 10 ms) and of the positive peak (here, around 20 ms), and the maximal slope of the curve in voltage over time (μV ms^−1^). Note that here the slope is represented with the tangent line at the inflection point.

#### Statistics

We run four Generalized Linear Mixed Models (GLMMs) on SPSS (IBM^©^ SPSS^©^ Statistics 27), to test the effect of condition (depth on anaesthesia: baseline, light sedation, deep sedation, and recovery) and stimulated brain region (frontal and posterior) over cortical excitability (amplitude, slope, positive, and negative latencies). The model took into consideration the attained DEX concentration as covariate, and the characteristics of the TMS pulse such as the Mean Induced Electric Field (V/m) and the distance of the electrode from the stimulation point in millimetres as random effects. Given that seven participants were still behaviourally responsive in the deep sedation condition (Ramsay score 3-4, UMSS 1-2 instead of the expected scores of 6 and 4, respectively), responsiveness at any condition was considered in the model as covariate (responsive vs unresponsive). Pairwise comparisons between conditions were performed with Bonferroni-adjusted two-tailed t-tests. We considered amplitude as the primary endpoint [P_critical_ = 0.05/(2 locations × 4 conditions) = 0.006], and slope, positive and negative latencies as secondary endpoints [P_critical_ = 0.05].

## Results

We modulated drug concentration to induce different conditions (sedation depth), which lead to different behavioural responses. The attained concentrations, as measured in the plasma for each condition were (mean ± SD, in ng mL^−1^): baseline:0 ± 0; light sedation: 1.37 ± 0.47; deep sedation: 3.41 ± 0.778; recovery: 2.71 ± 0.47. The UMSS score was (median, range): baseline: 0, [0 0]; light sedation: 2, [1 3]; deep sedation: 4, [2 6]; recovery: 2, [1 4]. Ramsay (median, range): baseline: 2, [2 2]; light sedation: 3, [3 4]; deep sedation: 6, [3 6]; recovery: 3, [2 5]. Interestingly, 7 out of 20 subjects were still responsive in deep sedation. As said before, we did not want to exceed our theoretical security threshold of a 2.5 ng mL^−1^ theoretical target to ensure the safety of our subjects. For more information about the participants, see **Table 1** and Appendix (**Table A1**).

According to our hypothesis, we found that condition (depth of sedation) modulated cortical excitability (see **Figure 3**). As shown in **Table 2**, and according to the GLMM models, there was a significant interaction between depth of sedation condition and stimulation location for amplitude [F_(3, 149)_ = 4.594, P = 0.004] and slope [F_(3, 149)_ = 4.009, P = 0.009], but not for the negative or positive peak latencies. Post hoc analysis (**Table 3**) showed that baseline amplitude in the frontal cortex was significantly different to the one in light sedation (Adjusted P < 0.0001), deep sedation (Adjusted P = 0.003) and recovery (Adjusted P < 0.001), while the slope in the frontal cortex was different in all pairwise contrasts (Adjusted P < 0.023), except for the deep sedation and recovery contrast (Adjusted P = 0.258). Slope and amplitude had the highest mean value in light sedation. These differences were not seen in the parietal region. Depth of sedation (**Table 2**) had an effect on positive peak latency [F_(3, 149)_ = 2.807, P = 0.042], but not on the latency of the negative peak [F_(3, 149)_ = 0.132, P = 0.22]. Irrespective of region, positive peak latency was significantly longer at light sedation than at baseline (Adjusted P = 0.030) (**Table 3**). The stimulated region (frontal vs. parietal) had an effect on negative peak latency [F_(1, 149)_ = 10.498, P = 0.001], meaning that it was globally significantly longer in the parietal region than in the frontal one, but no effect over the positive peak latency [F_(1, 149)_ = 1.234, P = 0.268]. Responsiveness to command had no effect on studied parameters (**Table 3**). **Figure 4** shows the effect of conditions and brain regions over cortical excitability. For more detailed information about the values of cortical excitability, see Appendix (**Table A2–A3**). We also removed the two subjects who showed the strongest effects (z-score>3) to test robustness of our findings and had virtually the same results with just small variations (see Appendix).

**Figure 3:**
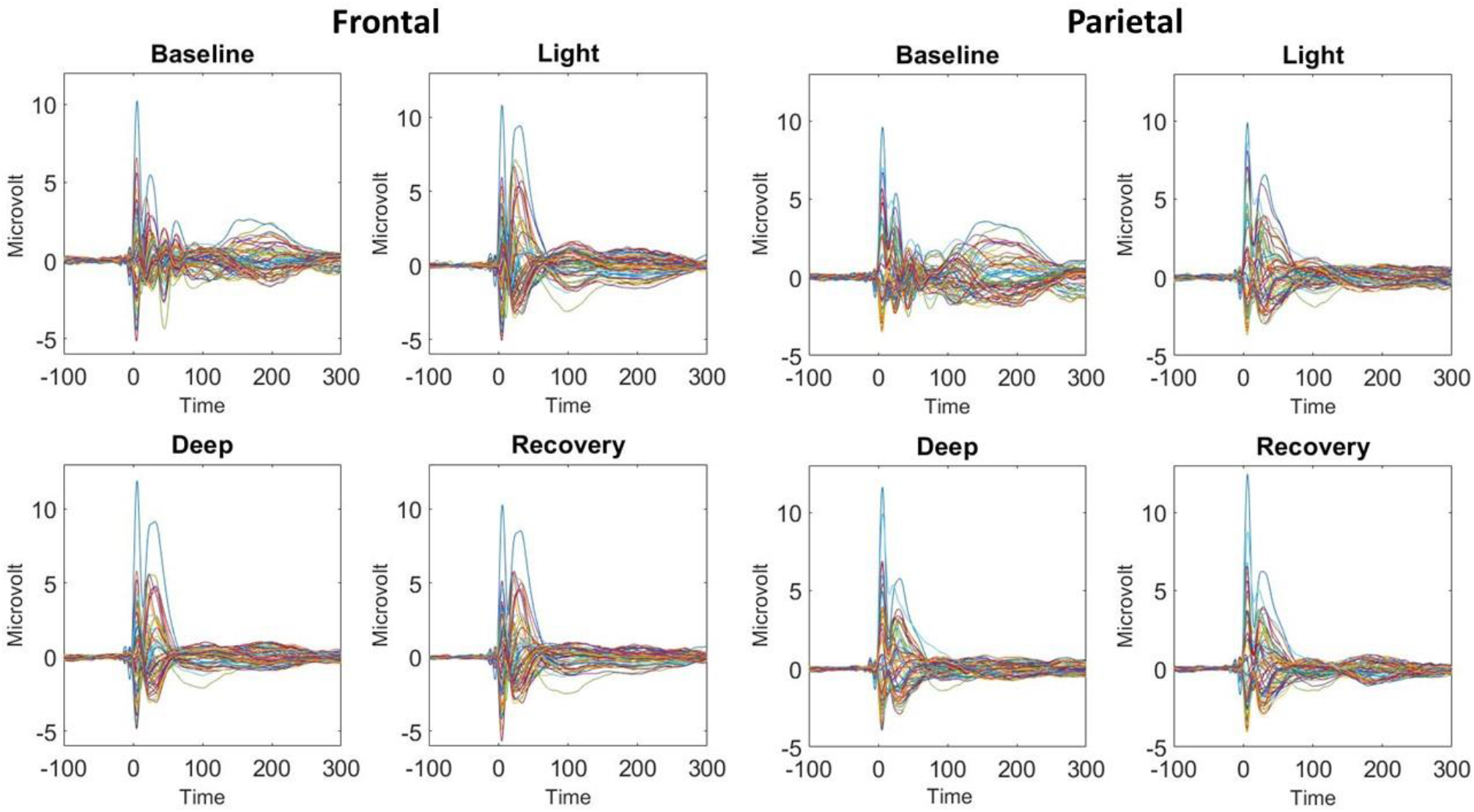
Grand average of the TMS-Evoked Potentials (TEPs) for all the subjects, divided by region (Frontal and Parietal) and the depth of sedation (baseline, light sedation, deep sedation, recovery).

**Figure 4:**
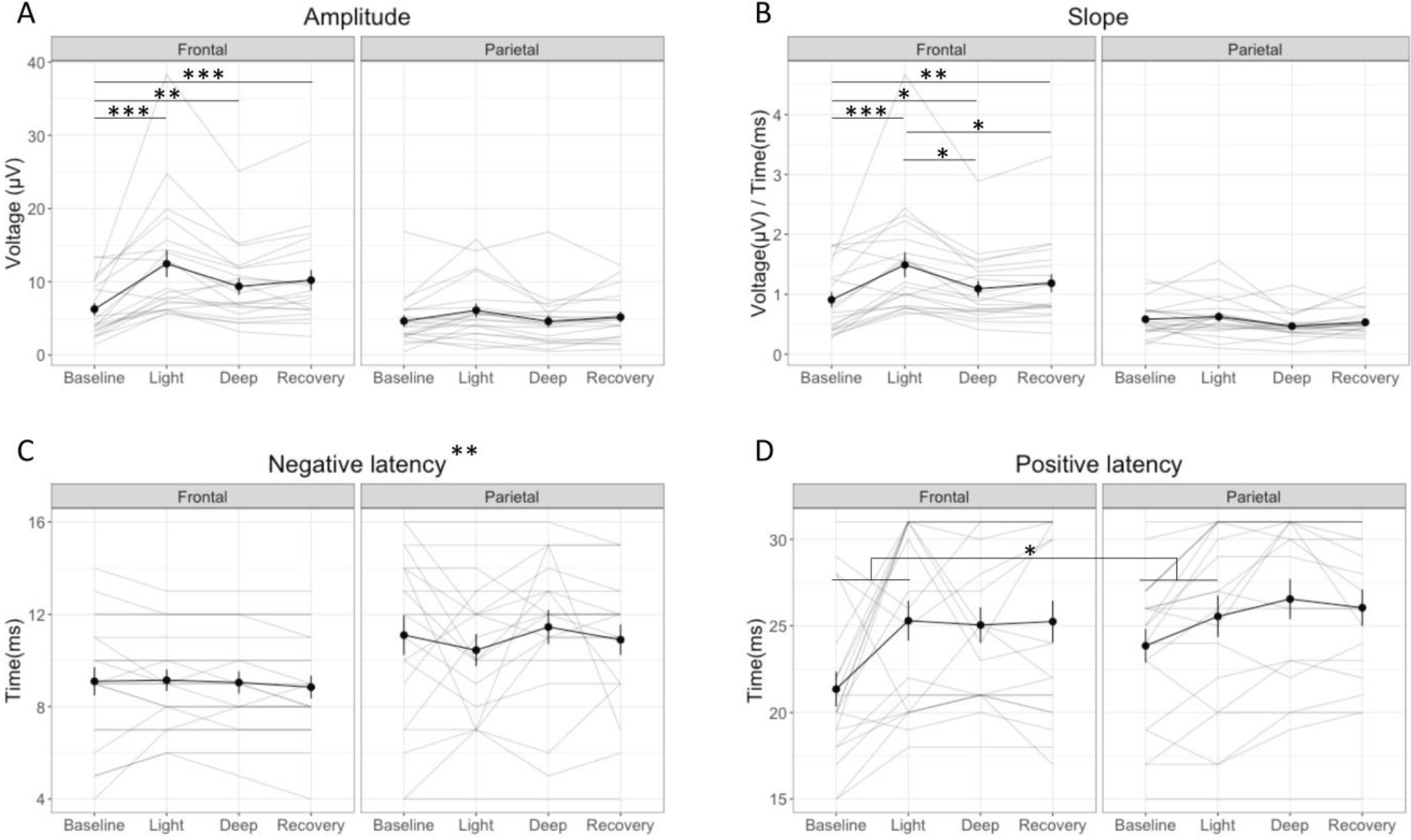
Averaged (black line) and individual results (grey line) of cortical excitability measurements (amplitude (A), slope (B), latency of negative peak (C) and latency of positive peak (D)) for the four conditions (baseline, light sedation, deep sedation, recovery). Each condition is divided according to the region (frontal vs. parietal). Error bars correspond to the standard error of the mean (SEM). The biggest change in amplitude and slope appears in the frontal cortex during light sedation in comparison to the other three conditions. For the image without the subjects who displayed the strongest effect, see Appendix (**Figure A1**). Legend: * = p<0.05, ** = p<0.01; *** = p<0.001

**Table 2.**
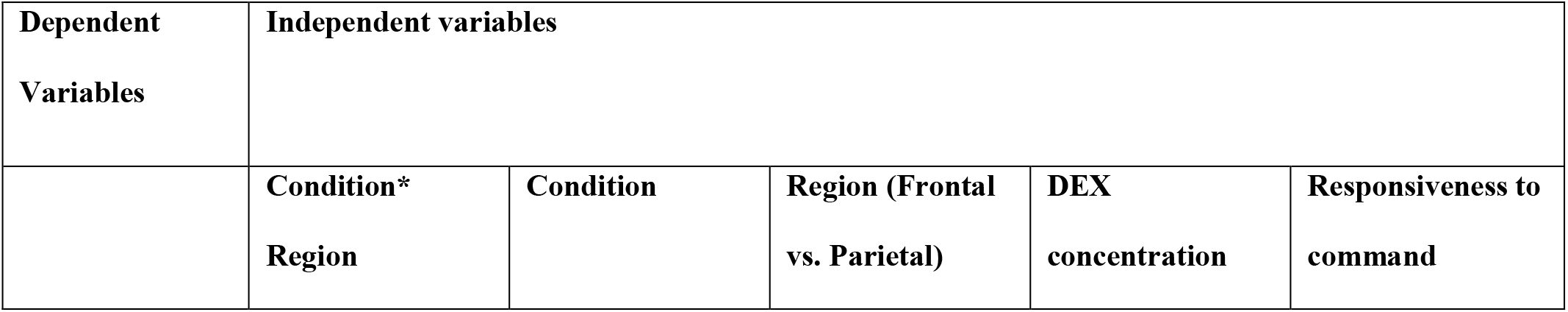
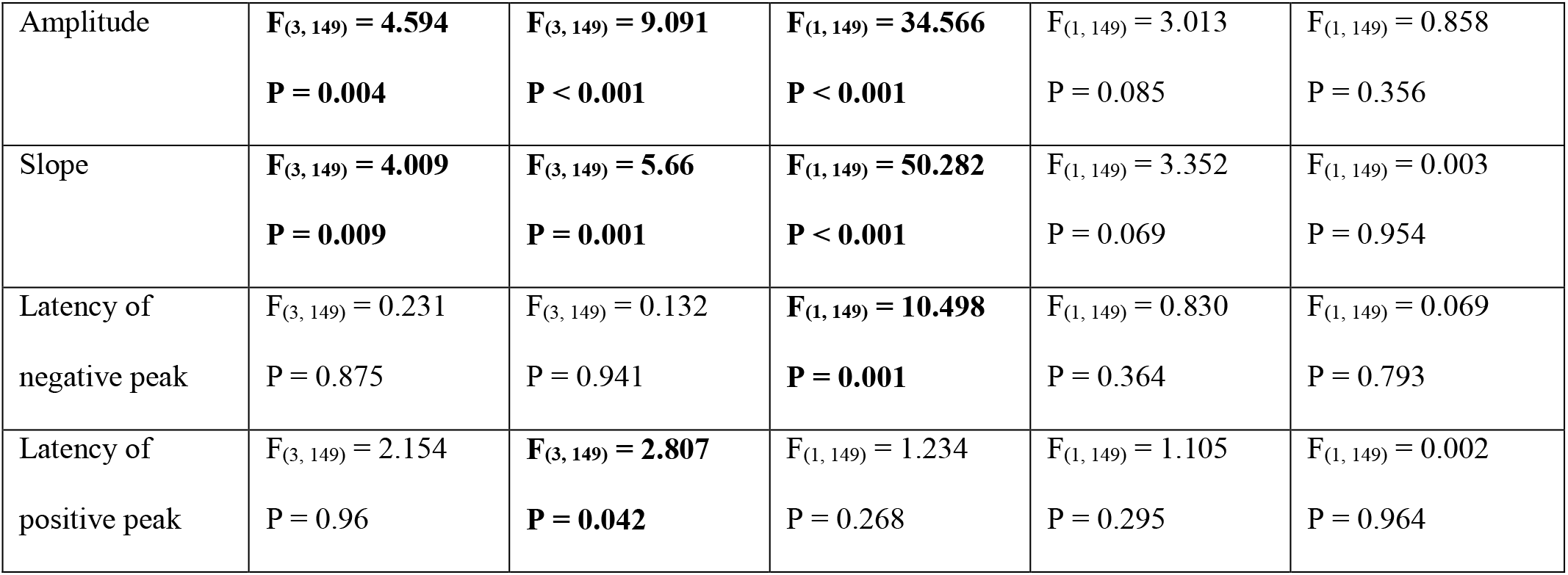
Results of the Generalized Linear Mixed Model (GLMM) on the modulation of cortical excitability. We took into consideration the condition (baseline, light sedation, deep sedation and recovery), the stimulated region (frontal vs. parietal), their interaction, the DEX concentration in the blood, and whether subjects were responsive in deep sedation. Significant effects in **bold**. Mean values and standard deviation of each measurement for condition are reported in the Appendix (**Table A2**).

**Table 3.**
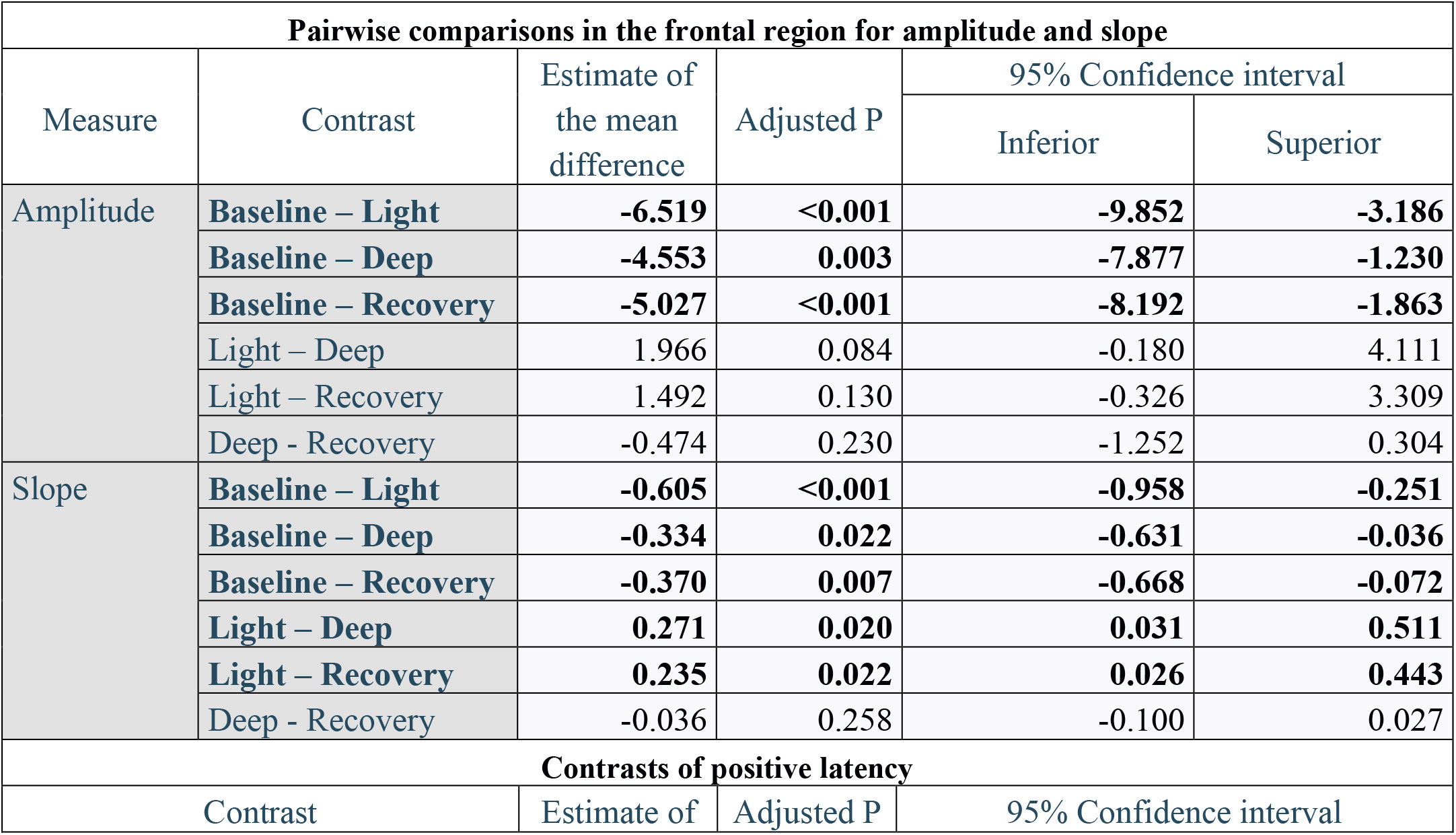
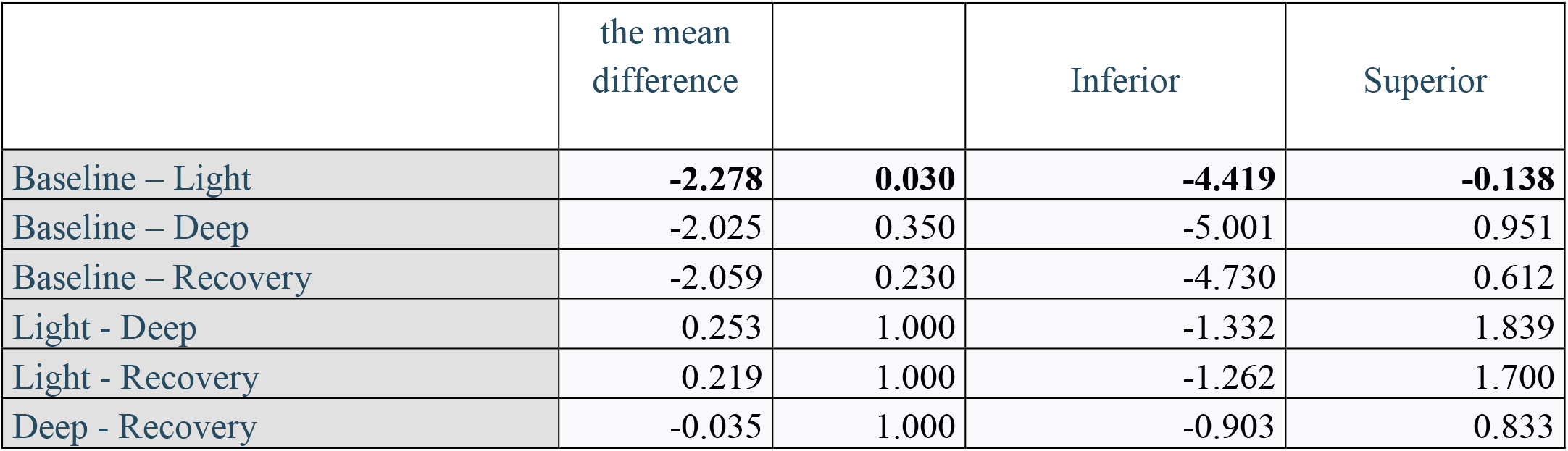
Post-hoc comparison for amplitude and slope in the frontal cortex, and for positive latency over the depth of sedation conditions. Significant comparisons are represented in **bold**. Since amplitude is our main endpoint, its P_critical_ is set to 0.006, while for slope and positive latency P_critical_ is 0.05.

Covariates as mean induced electric field, which summarize TMS pulse characteristics, and the distance of the electrode from which the TEP was taken, had a significant effect over cortical excitability. These effects are negligible and not informative for our purpose, as they were constant across conditions and had a smaller effect size compared to the effects of condition or brain region. They are reported in the Appendix (**Table A4**).

## Discussion

In the current study, we measured changes in cortical excitability as a function of the depth of DEX sedation in 20 healthy subjects, taking into consideration four conditions (baseline, light sedation, deep sedation, and recovery). Cortical excitability at the sensor level has been reported to increase during unconscious states, such as deep NREM sleep or disorder of consciousness like the unresponsive wakefulness syndrome.^21 26^ Thus, we expected cortical excitability to increase proportionally with the depth of sedation, being maximum during the deep sedation. According to our hypothesis, the condition had a strong effect on amplitude and slope. Interestingly, the effect was only present in the frontal cortex, and in contrast to our expectations, was not higher in the deep sedation compared to the light sedation, when subjects were drowsy but still able to respond to a command. To our knowledge, this is the first time that a non-linear evolution of cortical excitability is described under the action of an anaesthetic agent, in a region-specific manner. It is important to remark here that our definition of cortical excitability is purely operational in this context, in that it refers to the amplitude/slope of early TEPs, rather than to the nature of the underlying neuronal events. In fact, various mechanisms may account for the enhancement of early TEPs in DEX, including a stronger driving force in hyperpolarized postsynaptic neurons^27^, an increased discharge synchrony of cortical populations^28^, a reduction in synaptic depression^29 30^, and thalamic bursting triggered by the TMS-induced corticothalamic volley.

The increase of cortical excitability recorded in the frontal cortex is in line with two recent works about spontaneous conscious transition and TEPs.^22 31^ In the first one, cortical excitability of the motor cortex increased during drowsiness, but did not change in unresponsiveness, when participants were allowed to drift towards sleep during a detection task.^31^ In the second one, cortical excitability in the premotor cortex transiently increases during lapses of attention in a continuous attentive task after the usual bedtime, compared to no-lapses periods.^22^ These evidences support the idea that the reactivity of the frontal cortex is specifically altered in drowsy conditions where subjects might have impaired (but evident) behavioural responsiveness, as the one here described during light sedation. In congruence with this view, we observed higher amplitude in recovery (that is, after regaining response to command) compared to baseline. Arguably, during recovery, participants were in a state that was closer to light sedation than to baseline, being still drowsy. In other words, our results extend with a chemical manipulation what was previously reported in natural settings. It is however possible that these effects are specific to the sleep-like modulation of DEX and might not extend to other anaesthetics that are not α_2_-adrenergic agonists. If it is an effect of the sedation *per se*, cortical excitability might be a novel index of drowsiness and sedation, whose neural mechanisms should be investigated. However, the absence of the effect in the parietal cortex is a peculiar observation, as there are several reports that highlight the role of parietal regions for the emergence of consciousness.^32^ Future research should address this phenomenon in more detail.

Drug modulations of TMS-evoked responses are of paramount importance to depict the underlying neural dynamics of the compound, and to bridge neurochemical pathways, brain mechanisms, and behaviour. If there are a number of studies that inferred cortical excitability with TMS looking at changes in the resting state motor thresholds^33 34^, just a few observed TEPs.^31 35^ One issue is that TEPs (and in general evoked-responses) change from region to region,^36–38^ as proven by the effect of the region over the negative latency (see **Figure 4**). This is relevant, as TEPs might be modulated not only by the depth of sedation *per se*, but by the changes of the oscillatory activity (in a power-^38^ or a phase-dependent^39^ manner). In fact, the depth of sedation causes spectral modification, in particular within the beta frequency band, which predicts responsiveness under anaesthesia^17^ and wakefulness^40^, and the alpha and delta band, which are modulated by DEX concentration and state of consciousness.^16^ As shown in a recent work, alpha and beta activity in Rhesus macaques after DEX anaesthesia differs between loss of consciousness, recovery of consciousness, and the recovery of the task performance at pre-anaesthesia level.^41^ Future investigations should pinpoint in finer details what is the relationship between responsiveness, natural oscillation, and TEPs.

The current study presents strong effects on cortical excitability, with the same trend present in almost all the subjects we recorded. However, there are still some relevant limitations. First, we used a behavioural assessment for inferring consciousness. The absence of behavioural responses does not always coincide with unconsciousness^42^ and subjects may have relatively preserved higher order cognitive processes (i.e., semantics) during the loss of responsiveness due to DEX.^14^ In other words, one could say that consciousness assessment should be refined with bedside neurophysiological measurements. Nevertheless, we are confident that our participants were in a deep sedation even if some were responsive, considered that the DEX concentration in the blood was very high. Still, the reason why some subjects were still responsive in deep sedation while others were not is not clear. This may have something to do with the accuracy of the model we used for target-controlled infusion, and/or to inter-individual variability in the sensitivity to DEX action. The mechanisms that lead to responsiveness to a certain drug should be approached in a systematic way. Given that brain dynamics^43^ and spectral power^16^ change after DEX administration in dose-dependent fashion, different patterns and biomarkers could be used to predict responsiveness. Here, as shown in **Table 1**, responsiveness in deep sedation had no effect over cortical excitability (P > 0.35), so probably it cannot be used to predict responsiveness as it is. Another possible problem is that we did not randomize the order of light and deep sedation. We cannot exclude that the high excitability of light sedation is driven just by an order effect. Other studies with randomization could ensure that this not the case. Additionally, we did not have any free recall after the sessions. This could have helped to understand the phenomenological status of subjects, even when they did not show any kind of response to verbal command. Finally, as previously mentioned, a comparison with other drugs might show the extent of our results and elucidate possible underlying dynamics.

This is relevant to comprehend which neuropathways are important in changing cortical excitability and what is its links to sedation in a drug (in)dependent-manner.

In conclusion, we provide here the first evidence of non-linear evolution of cortical excitability after DEX intake, as indexed by the first component (0-30 ms) of the TEP at the closest electrode to the stimulation hotspot. We demonstrated that DEX sedation increases local cortical excitability in a region-specific manner, but do not differs between sedation level. In particular, we had no difference in cortical excitability between light sedation, deep sedation and recovery. This is in line with recent findings that describe abnormal high cortical excitability during drowsiness in natural settings. Interestingly, the effect was present only in the frontal cortex, and not in the parietal one. These results foster new questions for possible investigations about the nature of sedation and drowsiness that will result in a deeper understanding of cortical dynamics during anaesthesia.

## Authors’ contribution

OB, VB, SL and RS designed the study. OB, SW, MK, JS, AV, and VB collected the data. PC analysed the data. Data interpretation was performed by all authors. PC drafted the article with the help of OG and VB. All authors revised it critically for important intellectual content and gave final approval of the revised manuscript.

## Acknowledgements

We thank all the volunteers who participated in our studies, Gilles Vandewalle for vital support in the implementation of TMS-hdEEG excitability computation, Mario Rosanova, Simone Sarasso, Matteo Fecchio, and Renzo Comolatti for valuable discussions on DEX effects and TMS-hdEEG interpretation. We finally thank Orion Pharma for the quantification of DEX concentration in the blood.

## Appendix

## Methods – Blood sampling and dexmedetomidine quantification

Sample preparation was performed using solid phase extraction (SPE). Aliquots of 250 μl of plasma were mixed with 675 μl of 0.1% formic acid in water and 75 μl of internal standard solution (medetomidine-d3, 1 ng/ml). Samples were then extracted with Sep-Pak® tC18 100 mg 96-Well Plates (Waters Corporation, Milford, MA, USA) using an Oasis 96-well plate extraction manifold (Waters). The evaporation residue was dissolved in 100 μl of a solution containing 30 % methanol and 70 % water.

The HPLC-MS/MS system consisted of an Agilent 1200 HPLC instrument (Agilent, Santa Clara, California, USA) and an AB Sciex QTrap4000 triple quadrupole mass spectrometer (AB Sciex LLC, Concord, ON, Canada). Separations were performed with Gemini C_18_ analytical column (150 × 2.0 mm, particle size 5 μm; Phenomenex, Torrance, CA, USA)) coupled with Gemini C_18_ precolumn (Phenomenex). The column oven temperature was + 28°C. The mobile phase consisted of two eluents: A was 0.1 % formic acid in water and B was 0.1 % formic acid in methanol. The HPLC gradient began and was held for 1 min at 10 % of B and was then ramped to reach 95 % at 7 min. Then B was decreased back to 10 % in 0.5 min and held there for 3.5 min. The run time was 11 min with a flow rate of 0.3 ml/min. Sample injection volume was 20 μl. The retention time of dexmedetomidine and medetomidine-d3 was approximately 5.5 min.

Mass spectrometric detection was carried out using positive Turbo Ion Spray (TIS) ionisation and multiple reaction monitoring (MRM) mode. The ion source temperature was +500°C. The nebulizer gas (Gas 1) and turbo gas (Gas 2) settings were 50. Curtain gas (nitrogen) was set to 16. The TIS voltage setting was 5000 V. The selected reactions were as follows: for dexmedetomidine, the precursor ion – fragment ion pair was m/z 201.2 - m/z 95.05, and for the deuterated internal standard it was m/z 204.2 – m/z 98.05. The dwell time was 300 ms for both ion transitions. The declustering potential was 55 V for both molecules. Entrance potential was 10 V. Collision energy was 22 V. Collision cell exit potentials were 7 V and 6 V.

The calculations for the quantification were based on peak area ratios of the analyse and the internal standard. The chromatograms were analysed and processed using AB Sciex software (Analyst® version 1.6.3). The standard curves were generated using linear regression with 1/x^2^ weighting.

## Methods – Descriptive statistics

**Table A1:** Demographics of participants, with age, weight, handedness and whether they were still responsive to verbal command with high doses of DEX.

| Subject | Age | Sex | Height (cm) | Weight (kg) | BMI | Handedness | Responsive in deep sedation |
| --- | --- | --- | --- | --- | --- | --- | --- |
| **1** | 21 | Male | 184 | 95 | 28.06 | Right | False |
| **2** | 22 | Female | 163 | 64 | 24.09 | Right | True |
| **3** | 26 | Male | 185 | 66 | 19.28 | Right | False |
| **4** | 24 | Male | 179 | 66 | 20.60 | Right | False |
| **5** | 25 | Female | 164 | 62 | 23.05 | Right | False |
| **6** | 23 | Female | 159 | 58 | 22.94 | Right | False |
| **7** | 23 | Male | 182 | 81 | 24.45 | Right | False |
| **8** | 28 | Female | 170 | 68 | 23.53 | Right | True |
| **9** | 24 | Male | 180 | 98 | 30.25 | Right | True |
| 10 | 27 | Male | 179 | 77 | 24.03 | Right | False |
| 11 | 24 | Male | 178 | 63 | 19.88 | Right | False |
| 12 | 24 | Female | 172 | 63 | 21.30 | Right | True |
| 13 | 22 | Female | 169 | 55 | 19.26 | Right | False |
| 14 | 19 | Male | 183 | 63 | 18.81 | Right | False |
| 15 | 24 | Female | 177 | 90 | 28.73 | Left | True |
| 16 | 26 | Male | 169 | 55 | 19.26 | Left | False |
| 17 | 19 | Female | 162 | 53 | 20.20 | Right | False |
| 18 | 24 | Male | 180 | 79 | 24.38 | Left | True |
| 19 | 27 | Male | 177 | 85 | 27.13 | Left | True |
| 20 | 25 | Female | 161 | 59 | 22.76 | Right | False |

**Table A2:** Mean and standard deviation (mean ± SD) of cortical excitability measurement for each condition (baseline, light sedation, deep sedation, recovery), divided for region (frontal and parietal).

| Measurements | Region | Descriptive statistics |  |  |  |
| --- | --- | --- | --- | --- | --- |
|  |  | Baseline | Light sedation | Deep sedation | Recovery |
| Amplitude | Frontal | 6.27 ± 3.29 | 12.46 ± 6.37 | 9.37 ± 3.36 | 10.22 ± 4.68 |
| | Parietal | $4.64 \pm 4.12$ | $6.10 \pm 4.14$ | $4.61 \pm 3.84$ | $5.18 \pm 3.36$ |
| Slope | Frontal | $0.91 \pm 0.39$ | $1.49 \pm 0.69$ | $1.09 \pm 0.32$ | $1.18 \pm 0.42$ |
| | Parietal | $0.58 \pm 0.43$ | $0.62 \pm 0.42$ | $0.47 \pm 0.38$ | $0.53 \pm 0.35$ |
| Negative Latency | Frontal | $9.10 \pm 2.17$ | $9.15 \pm 1.69$ | $9.05 \pm 1.81$ | $8.85 \pm 1.90$ |
| | Parietal | $11.10 \pm 2.91$ | $10.45 \pm 2.21$ | $11.45 \pm 2.85$ | $10.90 \pm 2.24$ |
| Positive Latency | Frontal | $21.35 \pm 5.09$ | $25.30 \pm 4.33$ | $25.05 \pm 3.78$ | $25.25 \pm 4.58$ |
| | Parietal | $23.85 \pm 3.51$ | $25.55 \pm 4.26$ | $26.55 \pm 4.20$ | $26.05 \pm 3.55$ |

**Table A3:** Distance of the electrode (mm) from the stimulation hotspot and induced electrical field (V/m)

| Subject | Region | Electrode Distance<br>(mm) | Induced EF (V/m) |
| --- | --- | --- | --- |
| 1 | Frontal | 27.73 | 136.29 |
|  | Parietal | 54.64 | 118.08 |
| 2 | Frontal | 31.40 | 113.52 |
|  | Parietal | 42.94 | 147.47 |
| 3 | Frontal | 16.40 | 135.08 |
|  | Parietal | 54.97 | 143.18 |
| 4 | Frontal | 35.23 | 105.54 |
|  | Parietal | 47.42 | 134.18 |
| 5 | Frontal | 25.48 | 118.69 |
|  | Parietal | 50.57 | 146.72 |
| 6 | Frontal | 31.11 | 119.22 |
|  | Parietal | 31.06 | 132.39 |
| 7 | Frontal | 38.39 | 121.29 |
|  | Parietal | 37.66 | 134.15 |
| 8 | Frontal | 44.10 | 118.32 |
|  | Parietal | 38.54 | 104.93 |
| 9 | Frontal | 33.90 | 130.57 |
|  | Parietal | 54.97 | 139.21 |
| 10 | Frontal | 36.69 | 110.97 |
|  | Parietal | 65.40 | 98.95 |
| 11 | Frontal | 20.06 | 144.04 |
|  | Parietal | 57.84 | 164.70 |
| 12 | Frontal | 23.32 | 125.36 |
|  | Parietal | 52.70 | 145.11 |
| 13 | Frontal | 28.04 | 133.41 |
|  | Parietal | 63.73 | 139.82 |
| 14 | Frontal | 58.86 | 125.15 |
|  | Parietal | 51.62 | 130.96 |
| 15 | Frontal | 54.18 | 117.35 |
|  | Parietal | 45.06 | 136.24 |
| 16 | Frontal | 16.06 | 130.50 |
|  | Parietal | 58.15 | 119.71 |
| 17 | Frontal | 23.77 | 106.38 |
|  | Parietal | 26.41 | 139.41 |
| 18 | Frontal | 27.68 | 125.81 |
|  | Parietal | 27.30 | 138.20 |
| 19 | Frontal | 26.12 | 128.86 |
|  | Parietal | 31.97 | 131.52 |
| 20 | Frontal | 25.06 | 120.28 |
|  | Parietal | 55.15 | 144.03 |

## Results – Additional results of GLMM

**Table A4:** Results of induced electric field caused by the TMS, and the distance of the electrode from the hotspot, for the four measures of cortical excitability. In **bold**, significant results, and in *italics* tendencies. Note that the induced electric field is not significant for amplitude (P = 0.009) as it is the primary endpoint and we have corrected for multiple comparison (P_critical_ = 0.006). SPSS reported that induced electric field was redundant for the negative peak and gave no results for it.

| Dependent Variables | Independent variable |  |
| --- | --- | --- |
|  | Electric field (V/m) | Electrode Distance (mm) |
| Amplitude | $Z = 2.598$<br>$P = 0.009$ | $Z = 1.357$<br>$P = 0.175$ |
| Slope | **$Z = 4.048$**<br>**$P < 0.001$** | **$Z = 1.988$**<br>**$P = 0.047$** |
| Latency of negative peak* | $Z = /$<br>$P = /$ | $Z = 1.388$<br>$P = 0.165$ |
| Latency of positive peak | $Z = 0.602$<br>$P = 0.547$ | $Z = 0.864$<br>$P = 0.387$ |

## Results – GLMM without subjects with strong effect

To control the robustness of our results, we rerun the analysis excluding the subjects who showed the strongest results. As visible in **Figure 4,** two subjects presented relatively high amplitude (normalized amplitude > 3 standard deviations). The results without these two participants are virtually identical to the presented in the main text, showing a strong effect of condition over amplitude and slope, in particular in the frontal cortex. In **Figure A1**, we show the individual and average results by condition and cortical excitability parameters for all participants, while **Table A5** displays the significance of the results.

**Figure A1:**
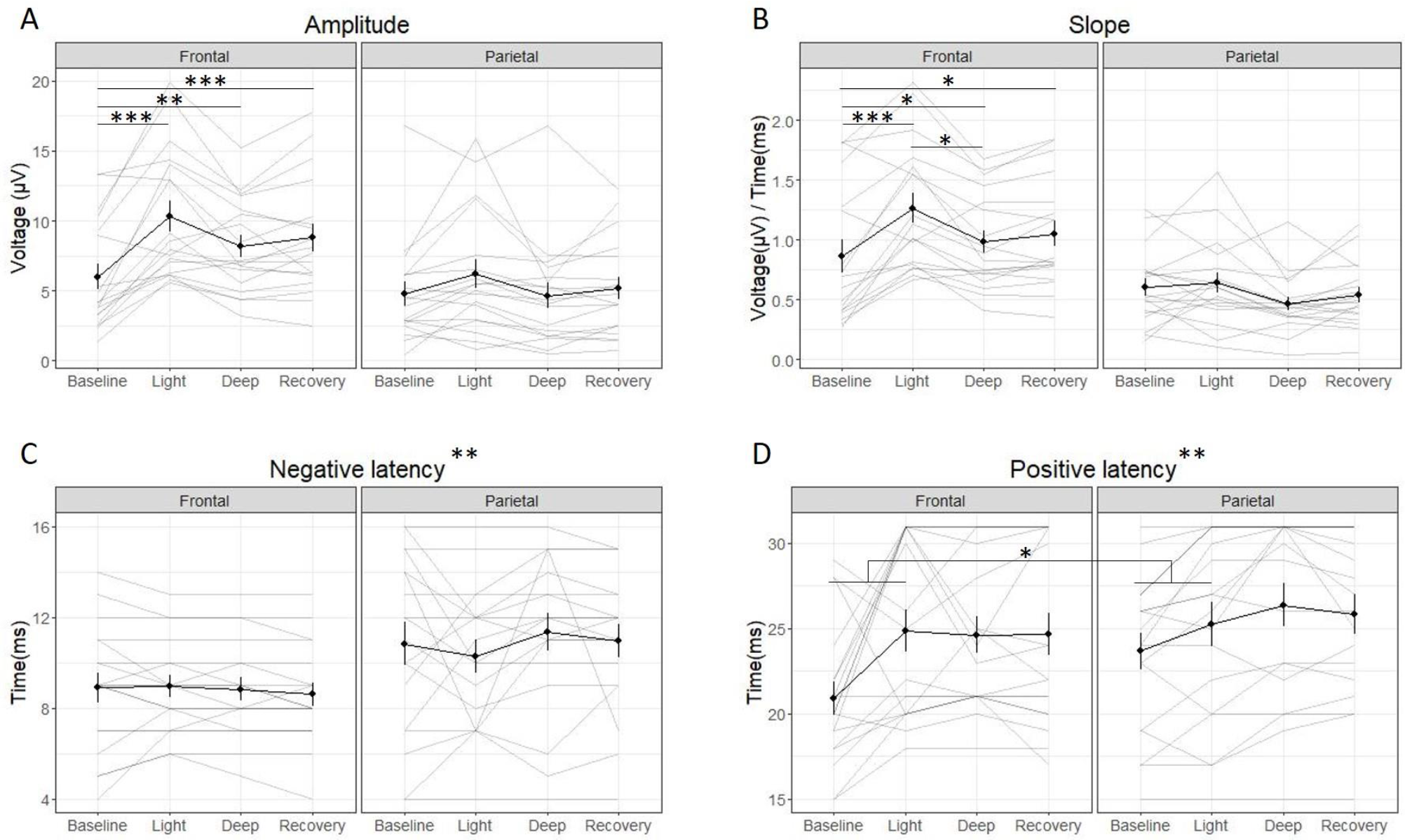
Averaged (black line) and individual results (grey line) of cortical excitability measurements (amplitude (A), slope (B), latency of negative peak (C) and latency of positive peak (D)) for the four conditions (baseline, light sedation, deep sedation, recovery). As Figure 4, divided in region (frontal vs. parietal), and with the standard error of the mean (SEM). The biggest change in the amplitude and slope appears in the frontal cortex during light sedation in comparison to the other three conditions. Legend: * = p<0.05, ** = p<0.01; *** = p<0.001

**Table A5.** Results of the Generalized Linear Mixed Model (GLMM) on the modulation of cortical excitability, without the subjects who had the strongest effect. Significant factors are in **bold**. For contrast and estimate of the contrasts of amplitude and slope in the frontal region, see **Table A6**.

| Dependent Variables | Independent variables |  |  |  |  |
| --- | --- | --- | --- | --- | --- |
|  | Condition | Region (Frontal vs Parietal) | Condition* Region | DEX concentration | Responsiveness in deep sedation |
| Amplitude | $F_{(3, 133)} = 5.319$<br>$P < 0.001$ | $F_{(1, 133)} = 9.352$<br>$P = 0.003$ | $F_{(3, 133)} = 3.513$<br>$P = 0.017$ | $F_{(1, 133)} = 3.709$<br>$P = 0.056$ | $F_{(1, 133)} = 1.817$<br>$P = 0.180$ |
| Slope | $F_{(3, 133)} = 6.552$<br>$P < 0.001$ | $F_{(1, 133)} = 18.948$<br>$P < 0.001$ | $F_{(3, 133)} = 3.716$<br>$P = 0.013$ | $F_{(1, 133)} = 4.460$<br>$P = 0.037$ | $F_{(1, 133)} = 0.052$<br>$P = 0.819$ |
| Latency of negative peak | $F_{(3, 133)} = 0.171$<br>$P = 0.916$ | **$F_{(1, 133)} = 10.555$**<br>**$P = 0.001$** | $F_{(3, 133)} = 0.329$<br>$P = 0.804$ | $F_{(1, 133)} = 0.952$<br>$P = 0.331$ | $F_{(1, 133)} = 0.056$<br>$P = 0.813$ |
| Latency of positive peak | **$F_{(3, 133)} = 2.725$**<br>**$P = 0.047$** | **$F_{(1, 133)} = 9.887$**<br>**$P = 0.002$** | $F_{(3, 133)} = 1.431$<br>$P = 0.237$ | $F_{(1, 133)} = 1.329$<br>$P = 0.251$ | $F_{(1, 133)} = 0.651$<br>$P = 0.421$ |

**Table A6:**
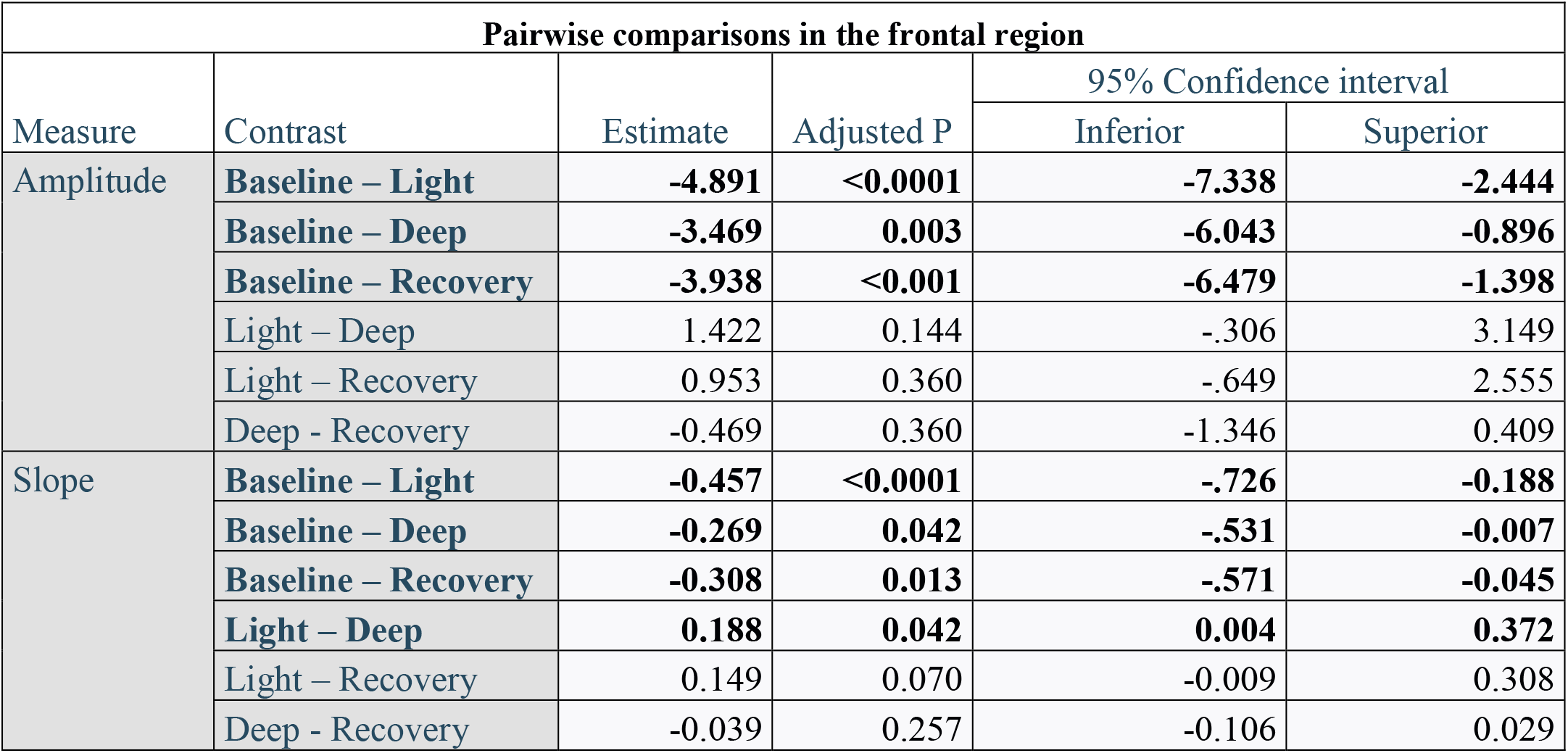
Post-hoc comparison for the amplitude and slope in the frontal cortex. Significant comparisons are represented in **bold**. Since amplitude is our main endpoint, its P_critical_ is set to 0.006, while for slope P_critical_ is 0.05.

Once these two participants were removed from the analyses, the general interpretation does not change: the effect of region is still present, with amplitude higher in frontal regions than in parietal regions. Critically, pairwise differences are exactly the same (as before, no significance of parietal cortex; compare **Table 3** and **Table A6**), so that it is just a region-specific effect, with an unexpected higher cortical excitability in light sedation. However, there are minor differences to what it is reported in the main text concerning the interaction between region and condition for the amplitude, the effect of DEX concentration for the slope, and the effect of the region over the positive latency. The two subjects who showed the strongest effect had a significant effect over the frontal cortex. If we observe instead the effect of DEX blood concentration over the slope, we can appreciate an effect that was not present before. However, given that the DEX concentration changes according to the condition, it is not surprising. Finally, we see here an effect of region to the positive latency. As said in the discussion, every region creates different evoked potential when stimulated, that thus creates a specific TEP for that region. So, even if beyond the scope of our paper, it is reasonable that the positive latency changes, as the negative one did.

**Table A7:** Results of induced electric field caused by the TMS, and the distance of the electrode from the hotspot, for the four measures of cortical excitability. In **bold**, significant results, and in *italics* tendencies. Note that the induced electric field is not significant for amplitude (P = 0.009) as it is the primary endpoint and we have corrected for multiple comparison (P_critical_ = 0.006). SPSS reported that induced electric field was redundant for the negative peak and gave no results for it.

| Dependent Variables | Independent variable |  |
| --- | --- | --- |
|  | Electric field (V/m) | Electrode Distance (mm) |
| Amplitude | Z = 1.595<br>P = 0.111 | **Z = 2.211**<br>**P = 0.027** |
| Slope | Z = 1.305<br>P = 0.192 | Z = 1.732<br>P = 0.083 |
| Latency of negative peak* | Z = /<br>P = / | Z = 1.122<br>P = 0.262 |
| Latency of positive peak | Z = 0.721<br>P = 0.471 | Z = 1.422<br>P = 0.155 |

